# Prior exposure to hypoxia alters DNA methylation patterns in the eastern oyster

**DOI:** 10.64898/2025.12.15.694390

**Authors:** Julia G. McDonough, Thomas J. Miller, Teresa W. Lee, Sarah C. Donelan, Sarah A. Gignoux-Wolfsohn

## Abstract

Environmentally induced epigenetic changes (e.g., DNA methylation) can alter genetic activity to help organisms adapt and respond to variable environments. While many studies have investigated DNA methylation as a response to a stressor at a single timepoint, less well-understood is how methylation may encode memory of past environments and influence the response to current environments (i.e., carryover effects). Oysters are an excellent natural system to study carryover effects due to their sessile nature, which may expose them to increased environmental variability. To better understand how methylation changes in response to a previous exposure of environmental stress, we conducted a fully factorial experiment exposing juvenile oysters to either control or hypoxic conditions at two timepoints separated by 60 days. After the second exposure, whole body tissue samples were collected and processed for methylRAD sequencing. Regardless of treatment, methylation was mostly found in exons. We found both the first and second exposure treatments contributed significantly to the observed variation in gene body methylation. Interestingly, oysters that were first exposed to hypoxia and later exposed to control conditions had methylation patterns that differed the most from any other condition. We found that differentially methylated genes identified in pairwise comparisons were mainly involved in the oxidative stress response, metabolism, and transcription. Together, these findings suggest that early life environments have a lasting impact on the epigenome and that the timing of stress elicits unique response strategies, which highlights potential targets of resilience for oysters.

## Introduction

Climate change is causing increased environmental variability (Pörtner et al. 2019), posing a challenge for organisms that have adapted to historical conditions and are now being repeatedly exposed to a wider range of stressors. Organisms can adapt and respond to these fluctuating stressors through environmentally induced epigenetic changes more quickly than by accumulating DNA mutations (Bollati and Baccarelli 2010; Eirin-Lopez and Putnam 2018). One of these epigenetic modifications, DNA methylation, occurs when a methyl group is added to a nucleotide in a DNA molecule, canonically on a cytosine that is next to a guanine (CpG) to form 5-methylcytosine (Mattei et al. 2022). CpG methylation can be added or removed in response to environmental changes (Head 2014), as observed in a wide variety of taxa including plants (Fan et al. 2013; Tang et al. 2018), invertebrates (Gupta and Nair 2021; 2025; Hawes et al. 2018; Venkataraman et al. 2020), and vertebrates (Bind et al. 2014; Yauk et al. 2008).

While DNA methylation is responsive to the environment, it can also be maintained long-term through mitotic divisions (Levenson and Sweatt 2006; D’Urso and Brickner 2014; Ming et al. 2021) and even sometimes across multiple generations (Fallet et al. 2020; Feiner et al. 2022; Liew et al. 2020). Methylation is therefore capable of encoding extended epigenetic memory about prior environmental conditions (D’Urso and Brickner 2014). The impacts of past environments on the response to current environments, whether physical, behavioral, or molecular, are often referred to as carryover effects (alternatively referenced as plasticity, parental effects, and legacy effects; O’Connor et al. 2014; West-Eberhard 2003). These carryover effects have been observed across developmental stages and different timepoints within a developmental stage (Hettinger et al. 2013). In some cases, carryover effects can be beneficial, when environments experienced in early life are a reliable indicator of the future environment and therefore prime the organism to be more stress tolerant upon repeated exposures (Kasumovic 2013). Priming of stressors during early life stages has been effective for increasing stress tolerance in geoduck clams (*Panopea generosa*) exposed to pCO_2_ (Gurr et al. 2022), sea anemones (*Nematostella vectensis*) exposed to acute heat stress (Glass et al. 2023), and juvenile eastern oysters (*Crassostrea virginica*) exposed to predation cues (Belgrad et al. 2021). Alternatively, carryover effects can be harmful and constrain later phenotypic plasticity. For instance, the proper development of claw asymmetries in juvenile lobsters relies on early life exposure to substrate (Latini et al. 2025). Once established, this morphological asymmetry is permanent and does not self-correct, even in cases when the claw is lost and subsequently regenerates.

Eastern oysters provide an excellent system in which to explore the epigenetic response to variable environments as they are relatively long-lived and sessile as adults. These life history traits mean they cannot move when environments become unfavorable and therefore individuals must respond to survive. Additionally, oysters live in the shallow brackish waters of estuaries that exposes them to numerous stressors on both a daily and seasonal basis, such as hypoxia (<2 mg/L O_2_), warming, and changes to salinity and pH. These stressors have been shown to influence oyster growth (Donelan et al. 2021), reproduction (Boulais et al. 2017), physiology (Jones et al. 2019), settlement (Stasse et al. 2022), and mortality(Stevens and Gobler 2018). Both the seasonal and daily fluctuations in hypoxia, especially in water tributaries of major estuaries such as the Chesapeake Bay, are increasing in both frequency and intensity (Breitburg et al. 2018) due to the interacting anthropogenic changes of cultural eutrophication (Tyler et al. 2009) and warming (Hinson et al. 2022). Understanding oysters’ response to these fluctuating stressors and their potential for adaptation is especially critical because they are ecosystem engineers that provide humans with multiple ecosystem services such as shoreline protection (Scyphers et al. 2011), improved water quality (Bricker et al. 2020), carbon sequestration (Fodrie et al. 2017; Parker and Bricker 2020), and contributions to the coastal economy via aquaculture and wild fisheries (Grabowski et al. 2012).

Studies exploring methylation as a molecular carryover effect or as a facilitator of carryover effects in marine invertebrates remain scarce. In abalone (*Haliotis discus hannai*), Dai and colleagues showed adults that experienced embryonic hypoxic stress had increased methylation levels and decreased oxygen consumption rates when exposed to a repeated acute hypoxic stress suggesting a higher tolerance for hypoxia may have been induced by methylation (Dai et al. 2024). Similarly, Dang and colleagues exposed larval Hong Kong oysters (*Crassostrea hongkongensis*) to low pH conditions followed by out planting juveniles in two field sites of different environmental stabilities. In the variable, unstable site they found that oysters pre-conditioned to low pH had higher survival and a different methylation profile than the control pH oysters, again suggesting stress tolerance may be influenced by methylation (Dang et al. 2023).

In the present study, we conducted a fully factorial experiment exposing juvenile oysters to control or hypoxic conditions at two timepoints separated by 60 days and collected whole-body tissue samples for methylRAD analysis to better understand how methylation changes in response to environmental stress. We have previously shown that an early exposure to warming, hypoxia, or both warming and hypoxia can lead to phenotypic carryover effects in the form of differences in growth of juvenile oysters (Donelan et al. 2021; 2023). In addition, numerous studies have demonstrated that oyster DNA methylation changes in response to environmental stressors such as warming (Wang, Li, Wang, Que, et al. 2021), acidification (Venkataraman et al. 2020), and salinity stress (Johnson et al. 2022). We therefore hypothesize that evidence of molecular carryover effects will be observed in the methylome of oysters exposed to hypoxic stress and that early hypoxic stress will induce differentially methylated genes. To test this, we first assessed the overall CpG methylation level and location within the *C. virginica* genome. We then identified patterns of gene-body methylation associated with the environmental treatment, including differentially methylated genes (DMGs) and associated adaptive biological processes between pairwise comparisons of our four treatment combinations. Understanding how multiple exposures to environmental stress, like hypoxia, affects the methylome of oysters contributes to our growing knowledge of how epigenetics may facilitate adaptive responses and potentially prime individuals for a future environment.

## Methods

### Experimental design for exposure to hypoxic stress

The oysters used in this experiment are a subset of those from Donelan et al., 2021 (see for full details of the experimental design), which had an additional treatment (warming). We only used oysters from the ambient temperature treatments here to focus on the effect of a single stressor (hypoxia). The control oysters in this paper are referred to as normoxic/ambient in Donelan et al., 2021.

Briefly, we used three- to four-month-old oysters (3-5 mm shell height) from Horn Point Hatchery in Cambridge, MD that were moved to flow through tanks at the Smithsonian Environmental Research Center (SERC) on the Rhode River in Chesapeake Bay. These oysters were acclimated for six days before entering Phase 1 of the experiment. In both Phase 1 and Phase 2, oysters were exposed to diel-cycling hypoxic stress or control (normoxic) conditions, creating four fully crossed treatment combinations. The target dissolved oxygen concentrations were 100% saturation for control and 0.5 mg/L or 8% saturation for hypoxia, which were successfully maintained throughout each Phase (see Donelan et al. 2021). Dissolved oxygen concentrations were manipulated in a 24-hour cycle for hypoxic conditions. Each cycle consisted of a 3-hour draw down from control levels to 0.5 mg/L, followed by a 4-hour plateau at the same concentration, then a 3-hour ramp up to control conditions where it remained for 14 hours. LabView software was used to manipulate water dissolved oxygen (detailed in Burrell et al. 2016). The water temperature and pH were maintained at ambient Rhode River conditions and did not differ between treatments (Donelan et al. 2021). The water conditions were continuously checked, and probes were calibrated with an external probe (Orion Star A326, Thermo Scientific, Waltham, Massachusetts, USA). For both Phase 1 and 2, dissolved oxygen cycled 5 times a week with a 14-hour light:10-hour dark photoperiod.

At the start of Phase 1, oysters were randomly selected and placed together in a perforated plastic container (n = 150 per container, N = 1,800), which was then placed into an aquarium tank. There were 12 total tanks, half that received control water and half that received hypoxic water. Oysters remained in Phase 1 (August 2018) for 18 days with 13 days of diel-cycling hypoxia, then removed and placed in common garden tanks in ambient conditions for 60 days until the start of Phase 2. At the start of Phase 2, half the oysters from each Phase 1 hypoxia treatment were placed into the same tanks as Phase 1 such that half were in control water and half were placed into hypoxic water to create our four fully crossed treatment combinations of control control (CC), control hypoxia (CH), hypoxia control (HC), and hypoxia hypoxia (HH) (where the first letter indicates the Phase 1 treatment and the second letter indicates the Phase 2 treatment). There were 48 oysters in each tank and oysters remained in Phase 2 (October - November 2018) for 18 days, with 14 days of diel-cycling hypoxia. Immediately following Phase 2, whole-body tissue samples were placed in molecular grade ethanol for preservation (n = 5 across all tank replicates for each of the four treatment combinations).

### DNA isolation and methylRAD library preparation

DNA from these oysters was extracted using the Qiagen DNeasy blood and tissue kit in 96 well plate format (Qiagen, Germantown, MD, USA) following manufacturers protocols. All sample concentrations were then normalized to 50 ng/ul. MethylRAD libraries were prepared following Dixon and Matz (2021) (a modified version of Wang et al. (2015). We digested the extracted DNA using the methylation-dependent endonuclease FspE1 (NEB, Ipswich, MA, USA). Digests were prepared with 0.4 units of FspE1 and the recommended amounts of enzyme activator and CutSmart buffer in a final volume of 15 μl. Digests were incubated at 37°C for 4 hours and then heated to 80°C for 20 minutes to deactivate the enzyme. Modified versions of the mdRAD 5ILL and mdRAD 3ILLBC1 adapters (with the addition of a standard 20 bp priming site for a second indexing PCR with Nextera indexes; see Pagenkopp Lohan et al. (2017)) were then ligated to the digested DNA in the following reaction: 10ul of digest, 0.2 μM mdRAD 5ILL adapter, 0.2 μM of the mdRAD 3ILLBC1 adapter, 800 units of T4 ligase (NEB, Ipswich, MA, USA), and 1mM ATP (included in ligase buffer). Ligations were incubated at 4°C overnight and then heat-inactivated at 65°C for 30 min. We then performed a PCR to add unique combinations of Nextera Illumina indexes in the following reaction: 4.5 μl water, 12.5 μl KAPA HiFi HotStart Ready Mix (Kapa Biosystems, Wilmington, MA, USA), 1 μl of 10 μM forward and reverse indexing primers, 6 μl of ligation. PCRs were run using the following program: 95 °C for 5 min, 12 rounds of 98 °C for 20 s, 60 °C for 45 s, 72 °C for 45 s, followed by extension at 72 °C for 5 min. PCR products were then cleaned using Ampure XP PCR cleanup beads in a 1.8:1 ratio of beads to PCR product. While we also did not see a single band at 32 bp (plus adapters) as in Wang et al. (2015), we did not see as big a smear as Groves and Matz (2021).

### Sequencing and sequence processing

Indexed ligations were sequenced on one lane of a HiSeq 2500 (paired end 150 bp reads). Raw methylRAD sequences were cleaned and quality controlled with TrimGalore! (Martin 2011; Krueger et al. 2023) (v0.6.5) to remove adapters (automatic detection) and low-quality reads (Phred score > 33) to obtain clean, paired end reads. Quality of the trimmed reads was assessed with FastQC (Andrews 2010). Only reads that were between 20 and 40 base pairs long were kept, since the expected product size from MethylRAD sequencing is 32 base pairs. Trimmed reads were filtered to only keep those with a methyl group in the middle (CCNGG, CCGG, GGCC, GGXCC, where N could be A, C, or T and X could be A, G, or T). Reads were then aligned to the NCBI RefSeq assembly for *C. virginica* (NCBI Accession GCF_002022765.2) using Bowtie2 (Langmead and Salzberg 2012) with the flags –very-sensitive and –local. The alignment rates for all samples were between 86-90%. The resulting SAM files were converted to BAM files, sorted by position, and indexed using samtools for downstream analysis (Danecek et al. 2021).

Sequencing produced a total of 802 million raw reads across 5 replicate samples per treatment combination (20 total samples). After trimming adapter sequences and barcodes, 108 million reads remained (13.5%), 23 million of which were paired correctly. Properly paired mapping efficiency averaged 70.9%, giving a final total read count of 13 million, with a mean of 675,000 for the 20 samples (Supplementary Table 1; NCBI Sequence Read Archive: BioProject accession number PRJNA1327452)

### Designating regions of interest

Various feature files were created for downstream analysis from the *C. virginica* genome annotation file (NCBI Accession GCF_002022765.2), based on methods outlined in Venkataraman et al. (2020). Exons, CDS, and genes were extracted from the GFF file and were converted to BED files using bedtools (Quinlan and Hall 2010). Untranslated regions (UTRs) of exons were identified by subtracting CDS from exons. Non-coding regions were identified by using complementBed from bedtools against exons. Introns were then identified using intersectBed from bedtools using genes and non-coding regions. Putative promoters were identified as 1KB upstream of the transcription start site. Hereafter, ‘gene bodies’ refer to the transcriptional region of a gene which includes exons, introns, and untranslated regions.

### Methylation of CpG dinucleotides

The observed sequencing depth for MethylRAD sequencing directly correlates to the degree of methylation. Methylation coverage of CpG dinucleotides for each sample was identified with bedtools multicov. For reliability, only CpG dinucleotides with at least 5 sequences were considered methylated. The BEDtools suite (Quinlan and Hall 2010) was used to determine the location of the methylated CpGs in relation to putative promoters, exons, introns, UTRs, and intergenic regions. Only CpGs that were methylated (>5 sequences) in the majority of replicates for each treatment combination were considered for this analysis. A chi-squared contingency test (chisq.test in R Version 4.3.2) was used to assess the association between genomic feature and methylation status of CpG dinucleotides. A two-way ANOVA (aov in R Version 4.3.2) was used to assess differences in CpG methylation level between treatments, and a post-hoc Tukey honestly significant difference (HSD; TukeyHSD in R Version 4.3.2) test was used to compare the means of CpG methylation between treatments.

### Differential methylation

A counts matrix of methylated sequences within gene regions was generated with featureCounts (Liao et al. 2014) using the GFF file (NCBI Accession GCF_002022765.2) and sorted BAM sequence files (GTF.featureType=”gene”, useMetaFeatures=TRUE, isPairedEnd=TRUE). All replicate samples had successful assigned alignments around 70%. To better understand how the timing of hypoxic stress influences DNA methylation, DESeq2 (Love et al. 2014) was used to identify differentially methylated genes (DMGs) in all possible pairwise comparisons (CC vs. HC, CH vs. HC, HH vs. HC, HH vs. CH, HH vs. CC, CC vs. CH). A log fold change normal shrinkage estimator was applied to DESeq results with the argument lfcThreshold=0.25 and type=”normal”. Shrinking log fold changes allows comparison of log fold changes across treatments and the ability to rank genes for downstream analysis. A gene was identified as differentially methylated if the absolute log fold change was greater than 0.5 and the adjusted p-value was less than 0.05.

Non-metric multidimensional scaling (NMDS) plots were used to visualize intragenic (only exons, introns, UTRs) methylation patterns of the four unique treatment combinations (CC, CH, HC, and HH). The DESeq2 results were normalized and transformed using variance stabilizing transformations (VST) using the blind dispersion estimation, which was all done within the DESeq2 package (Love et al. 2014). Differences in gene body methylation patterns among treatment combinations were tested using PERMANOVA (Anderson 2017) in the vegan package (Oksanen et al. 2009) in R. Volcano plots were used to visualize differential methylation data between each pairwise comparison. A Venn diagram was used to visualize the overlap of DMGs across pairwise comparisons with ggvenn (Yan 2023). Finally, gene ontology analysis was performed on the differentially methylated genes with GO-MWU, a rank-based gene ontology analysis (Wright et al. 2015). The generic GO subset was used to match GOslim terms to DMGs for broader categorization into biological processes (Ashburner et al. 2000; The Gene Ontology Consortium et al. 2023).

## Results

### CpG Methylation Level and Genomic Distribution

To better characterize methylation in the eastern oyster genome, we examined the distribution of methylated CpGs across six genomic features: exons, intergenic regions, introns, putative promoters (defined as 1KB upstream of the ORF) and untranslated regions. We found that the proportion of methylated CpGs in each feature category significantly differed from the distribution of all CpGs across categories regardless of treatment combination (Contingency test; χ2 > 66000 for each treatment combination vs. all CpGs, df = 5, P = 2.2e-16; Figure 1A). Notably, although only 9.93% of genomic CpGs are found in exons, nearly 40% of methylated CpGs are found in exons (averaged across treatments), suggesting CpGs in these regions are preferentially methylated. Conversely, methylated CpGs were not as abundant in intergenic regions (13.84%) and, to a lesser extent, putative promoters (2.46%) when compared to all CpGs (28.40% and 3.96%, respectively). Introns and untranslated regions of exons all have methylation proportions that are equivalent to the distribution of all CpG dinucleotides.

**Figure 1.**
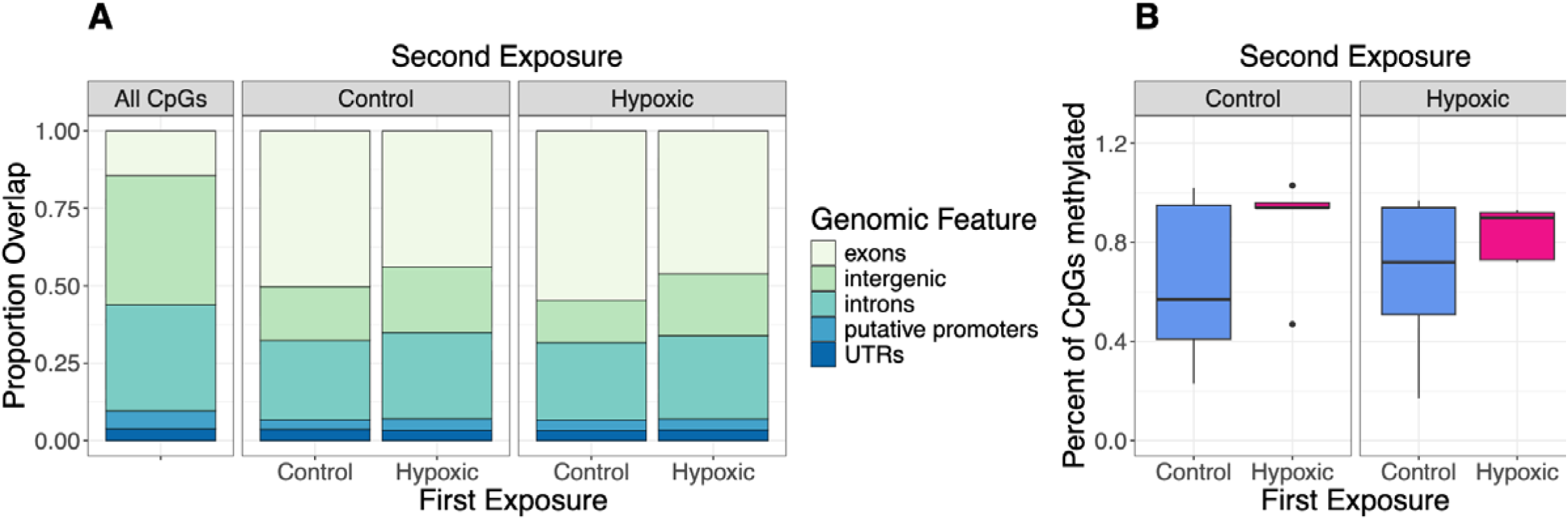
Methylation of CpG dinucleotides varies with genomic features and environmental exposure in oysters. (**A**) Proportion of all CpG dinucleotides (left panel) and methylated CpG dinucleotides across treatment combinations (two right panels) found in various genomic features. (**B**) Percent of all CpG dinucleotides that were methylated in oysters first exposed to either control (blue) or hypoxic (pink) conditions and second exposed to control (left) or hypoxic (right) conditions. The middle line of the box represents the median percent of CpGs that were methylated and the top and bottom edges of the box represent the first and third quartiles, respectively. Black points outside of the whiskers are outliers.

Consistent with prior observations in the eastern oyster, we found that CpG methylation is rare: only between 0.25% and 1.00% of CpG dinucleotides were methylated across all treatment combinations (Figure 1B). We found no significant differences in total levels of methylation based on either first (two-way ANOVA, F_1,16_ = 2.874, P = 0.109) or second (F_1,16_ = 0.000, P = 0.983) exposure, or the interaction (F_1,16_ = 0.049, P = 0.828), although the median number of methylated CpGs was slightly higher in oysters that experienced a first exposure to hypoxic stress (Figure 1B, pink), than in oysters who experienced a first exposure to control conditions (blue) regardless of the second exposure.

### Gene-body DNA Methylation Patterns

We next wanted to examine whether multiple exposures to hypoxic or control conditions influence the overall distribution and levels of methylation within gene bodies (only exons, introns, and UTRs). We found a significant effect of both the first (PERMANOVA, F_1,16_ = 1.364367, R2 = 0.068, P = 0.017) and second (F_1,16_ = 1.516458, R2 = 0.076, P = 0.004) exposures, which together explains roughly 15% of the total variation in gene body methylation, although the interaction of the first and second exposures was not significant (PERMANOVA, F_1,16_ = 1.169, R2 = 0.058, P = 0.088). We found oysters that were first exposed to control conditions have more dispersion in their methylation patterns, and were therefore more different than each other, when compared to oysters that were first exposed to hypoxic conditions (Figure 2A). When oysters were first exposed to control conditions, the second exposure to either treatment had almost no effect on overall methylation patterns. Conversely, when the first exposure was hypoxic, we saw a clear distinction between second exposure conditions (Figure 2A). This pattern becomes stronger when only considering differentially methylated genes (DMGs) identified in any pairwise comparison with DESeq2 (Figure 2B): 60% of the variation in gene body methylation of these genes can be explained by the significant effects of the first (PERMANOVA, F_1,16_ = 8.023, R2 = 0.200, P = 0.001) and second (F_1,16_ = 12.500, R2 = 0.311, P = 0.001) exposure treatments alone, as well as the interaction of the two treatments (F_1,16_ = 3.705, R2 = 0.092, P = 0.025). On the nMDS, samples cluster according to their first exposure, with clustering by the second exposure only occurring for those samples where the first exposure is hypoxic, indicative of molecular carryover effects (Figure 2B). Overall, HC oysters had methylation patterns that were the most different from all other treatment combinations, regardless of whether we considered all genes or DMGs.

**Figure 2.**
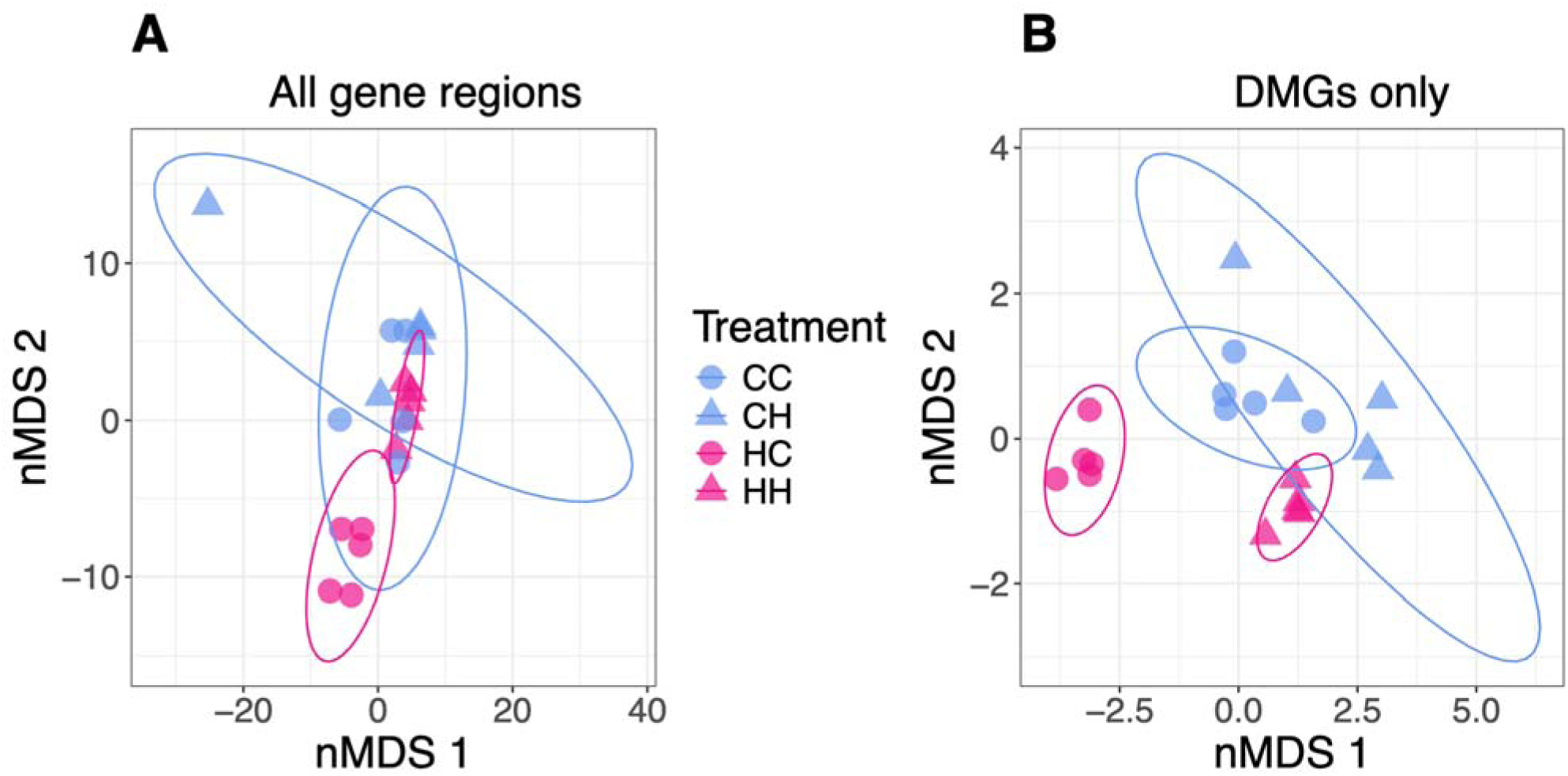
Differences in methylated gene regions are most pronounced in oysters first exposed to hypoxic and then control conditions compared to all other treatment combinations. nMDS plots of methylation patterns for (**A**) all genes and (**B**) significantly differentially methylated genes in any pairwise comparison for oysters with a first exposure to control (blue) or hypoxic (pink) conditions and a second exposure to control (circle) or hypoxic conditions (triangle).

### Differentially Methylated Genes

To see how a prior single exposure to stress affects DNA methylation after a period of recovery, we compared CC and HC oysters (Figure 3A). We identified 12 DMGs, one with more methylation in HC and 11 with more methylation in CC oysters. Of these 12 genes, two were involved in ubiquitin-dependent protein catabolic processes and one (GGT1) was involved in catalytic activity. We next examined how a single exposure to hypoxia only experienced later in life affects methylation by comparing CC to CH oysters (Figure 3B) – one DMG (MDE1) was different between these conditions and involved in metabolic processes. Taken with the results of the CC vs HC comparison, these results suggest that exposure to hypoxia earlier in life (16 weeks) has a larger effect on methylation than exposure later in life (27 weeks). Next, to explicitly look for differences in methylation based on the timing of initial hypoxic exposure we compared CH to HC oysters (Figure 3C) – this comparison had the largest difference, with 196 DMGs identified (60 genes more methylated in HC and 136 more methylated in CH), indicating that prior exposure to a stressor has an epigenetic effect that is distinct from the immediate response to the same stressor.

**Figure 3.**
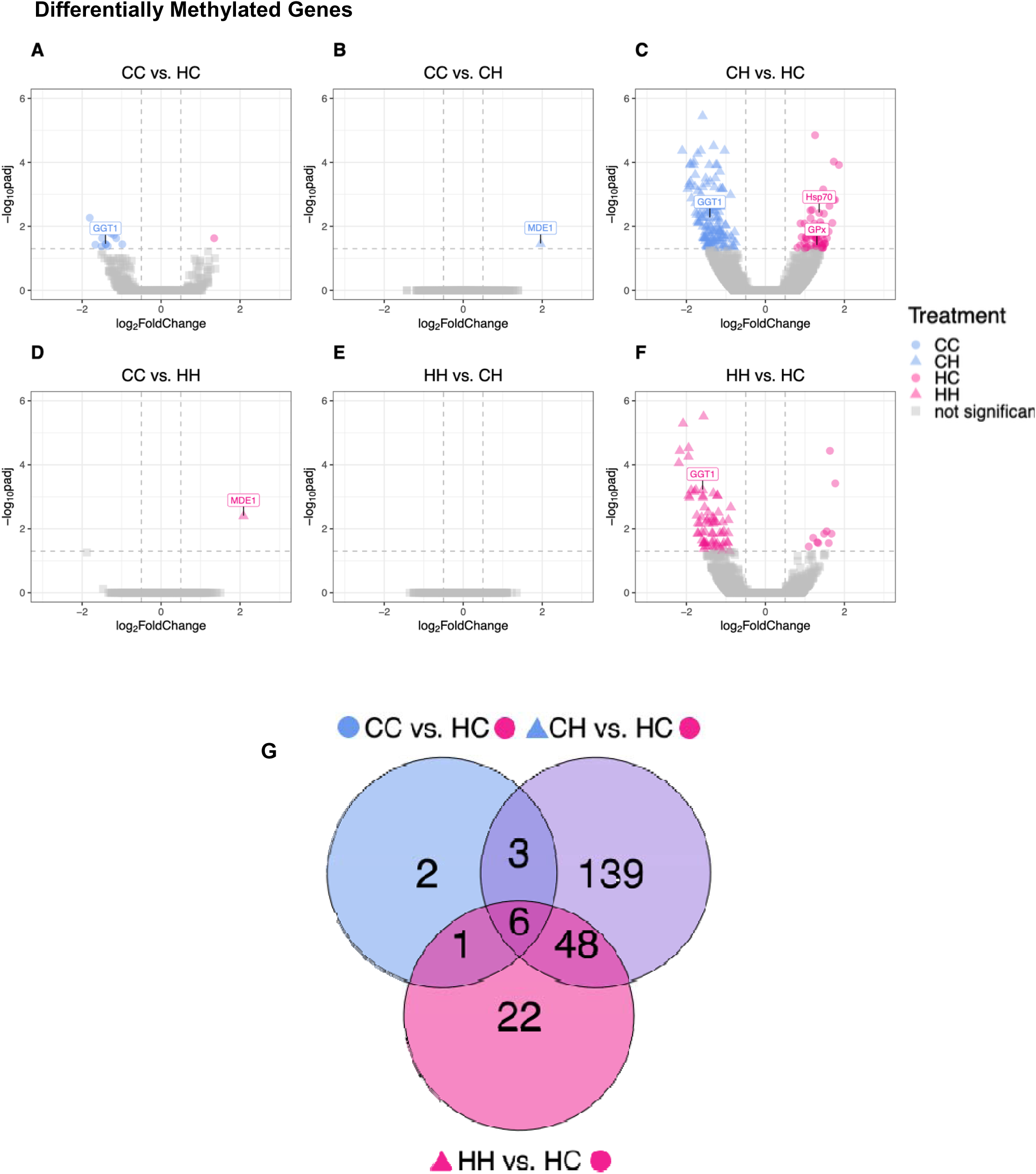
Timing of hypoxic exposure results in differentially methylated genes. (**A** to **F**) Volcano plots of DMGs from pairwise comparisons of hypoxic and control treatment combinations. Directionality of methylation is indicated by colored symbols: first exposure to control (blue) or hypoxia (pink) and second exposure to control (circles) or hypoxia (triangles). (**G**) Venn diagram of shared and unique significant DMGs in pairwise comparisons with HC oysters.

Unexpectedly, when we compared the oysters that experienced no stress (CC) to those that experienced the most stress (HH), we only found one gene (MDE1 again) that was differentially methylated (Figure 3D). To determine whether an initial exposure to stress affects the response to a later exposure, we next compared HH to CH oysters and identified no DMGs (Figure 3E), indicating that the more recent exposure to hypoxia overrode the effects of a previous exposure. Finally, we tested whether the initial stress exposure affects the subsequent response to stress by comparing HH to HC oysters and identified 77 DMGs (Figure 3F). Ten of these DMGs were more methylated in HH oysters and 67 were more methylated in HC oysters. These genes are involved in metabolic processes (6 DMGs), DNA-templated transcription, signaling, and protein transport (2 DMGs each).

To better understand the shared and unique responses to varying hypoxic stress, we compared DMGs across pairwise comparisons (Figure 3G). Consistent with our observations about overall gene body methylation patterns (Figure 2A), we saw that HC oysters had the largest number of DMGs compared to oysters in the other treatments. Six genes were consistently different between HC treated oysters and all three other treatments, indicating a shared response in HC oysters. Five of these genes are annotated, two were involved in cell signaling, one (GGT1) in catalytic activity, one in transcription, and one in developmental process (Figure 3G, Supplementary Table 2). We found the majority of shared DMGs (54 total) were between comparisons of HH and CH against HC (Figure 3G), where the genes are different because the oysters experienced a second exposure to hypoxia. These DMGs were mostly involved in metabolic processes (9 DMGs) and signaling (3 DMGs). Only one DMG was shared exclusively between the CC vs. HC and HH vs. HC comparisons (involved in exocytosis), suggesting that these comparisons exhibited distinct patterns of differential methylation. Three DMGs were shared between the CC vs. HC and CH vs. HC comparisons (unannotated), indicating a shared response to early control conditions. In comparisons outside of those against HC oysters, only one DMG was shared (MDE1) which had more methylation in oysters from the hypoxic treatments (HH and CH) than the control (CC) oysters.

We found that about 75% of DMGs (163 total) were unique to the comparison of CH with HC oysters (Figure 3G), indicating the response to hypoxic stress depends on the timing of when it is experienced. These DMGs were mostly involved in in signaling (10 DMGs) and metabolic processes (9 DMGs). We identified 22 DMGs that were unique to comparisons of HH and HC oysters, which suggests multiple stress exposures triggers an explicit methylated response. The majority of these genes were involved in metabolic processes (6 DMGs), DNA-templated transcription, signaling, and protein transport (2 DMGs each). Finally, only two DMGs were unique in comparisons examining the effect of an early hypoxic stress exposure after a period of recovery (CC and HC), and both genes were involved in ubiquitin-dependent protein catabolic processes.

In all pairwise comparisons, there were not any significantly enriched gene ontologies (Supplementary Table 3). Out of all identified DMGs in any pairwise comparison, 18.4% were uncharacterized and 41.1% did not match to any GO terms (Supplementary Table 2).

## Discussion

In the present study, we conducted a fully factorial experiment exposing juvenile oysters to control or hypoxic conditions at multiple timepoints and collected whole-body tissue samples to better understand how methylation changes in response to environmental stress. Our results showed that there are impacts of early exposure on methylation patterns and that these patterns can vary with the second exposure, supporting our hypothesis and providing evidence for molecular carryover effects. Additionally, many of the DMGs we identified are involved in metabolism, stress response, and transcription – future studies into the specific pathways affected by these DMGs will provide insight into the molecular basis of how oysters respond to hypoxic stress.

### Low global methylation is non-randomly distributed in *C. virginica*

We observed low levels of DNA methylation (between 0.2 and 1.2% of all CpG sites for individual oysters), in line with what has been reported for other marine invertebrates and significantly lower than what has been observed in vertebrates, which tend to have 30% to 80% of their genomes methylated (Klughammer et al. 2023; Dixon et al. 2016; Wang et al. 2014). However, our observed levels are lower than the ∼15% that has previously been reported in *C. virginica* (Venkataraman et al. 2020; 2024) and *C. gigas* (Venkataraman et al. 2022; Wang, Li, Wang, Que, et al. 2021). This discrepancy may stem from our use of MethylRAD-sequencing and stringent quality filtering that led to decreased sequencing depth in our analysis.

In vertebrates, DNA methylation is widely distributed across most genomic features and is well-known to primarily regulate gene activity (largely as a repressive mark). However, in invertebrates, the role of DNA methylation is less well understood, partially because patterns of methylation are more varied across taxonomic groups (Head 2014; Tweedie et al. 1997). For instance, DNA methylation is nonexistent in *Caenorhabditis elegans* (Simpson et al. 1986) and moderately abundant (roughly 30% of CpG dinucleotides) in *Hydra vulgaris* (Ying et al. 2022). These diverse patterns of methylation suggest that there is no single functional role for DNA methylation across all invertebrates.

In the eastern oyster, we found that the majority of methylation occurred within intragenic regions (primarily in exons), consistent with previous studies in the same species (Downey-Wall et al. 2020; Venkataraman et al. 2020), as well as studies in other species of oysters (Lim et al. 2021; Wang et al. 2014) and marine invertebrates (Dixon et al. 2016). Methylation of exons is likely to either directly regulate gene expression or modulate alternative splicing (Flores et al. 2012; Huang et al. 2016; Maunakea et al. 2010; Shukla et al. 2011). However, previous studies looking for correlations between DNA methylation and gene expression in oysters have found conflicting results. Positive associations between methylation and gene expression have been found in *C. gigas* in both gill tissue (Gavery and Roberts 2013) and male gametes (Olson and Roberts 2014) under control conditions. However, no correlation between methylation and gene expression patterns were found in *C. virginica* gill tissue when challenged with disease (*Perkinsus marinus*) (Johnson et al. 2020) or salinity stress (Johnson et al. 2022) and only a weak correlation in the mantle after exposure to ocean acidification (Downey-Wall et al. 2020). The relationship between DNA methylation and gene expression is known to vary across taxa. As previously mentioned, DNA methylation is usually negatively correlated with gene expression in vertebrates, while in some species like rice (*Oryza sativa*) and tunicates (*Ciona intestinalis*), moderately expressed genes are more likely to be methylated. In other taxa like anemones (*Nematostella vectensis*) and silk moths (*Bombyx mori*), methylation is found most in highly expressed genes (Zemach et al. 2010). Methylation can also influence alternative splicing, which may obscure observable relationships with gene expression (Lev Maor et al. 2015; Shayevitch et al. 2018). Given the conflicting potential roles of DNA methylation in gene regulation for this species, the lack of paired gene expression data in our current study limits our understanding of what biological effect these patterns are likely to have.

Similar to observations in *C. gigas* and other invertebrates, we found that methylated CpGs were less frequently found in putative promoters than gene bodies across all treatments (Venkataraman et al. 2020; Zemach et al. 2010). Promoters in both invertebrates and vertebrates are often comprised of transcription start sites and CpG islands, which are CpG-rich hypomethylated sequences (Angeloni and Bogdanovic 2021; Antequera and Bird 1993). Because of these elements, methylation of promoters often silences downstream gene expression (Lou et al. 2014). More recently, a context-dependent role for methylation of promoters has been suggested (Keller et al. 2016; Olson and Roberts 2014). Keller et al. (2016) demonstrated that methylated promoters only affect gene expression in the muscles of tunicates (*C. intestinalis*) when the adjacent gene body is also methylated. Further evidence of methylation influencing alternative promoter usage has similarly been shown in vertebrates (de Mendoza et al. 2022; Sarda et al. 2017). Despite low levels of observed promoter methylation in the present study, it is possible differential methylation of these regions may facilitate tissue-specific transcriptional and phenotypic differences.

### Multiple exposures to hypoxia alter methylation patterns

We found that DNA methylation was influenced by an initial exposure to hypoxic stress: juvenile oysters that experienced a first exposure to hypoxia had higher average levels of methylation compared to those in control conditions regardless of their second exposure (although this difference was not significant). Methylation remains consistently responsive to environmental conditions in oysters exposed to changes in pH (Downey-Wall et al. 2020; Lim et al. 2021; Venkataraman et al. 2020), salinity (Johnson et al. 2022; Zhang et al. 2017), temperature (Roberto et al. 2021; Wang, Li, Wang, Que, et al. 2021), or oxygenation (Wang et al. 2023; Wang, Li, Wang, Zhang, et al. 2021). Similar relationships between environmental stress and increased DNA methylation have been observed in other species, including the stony coral *Pocillopora damicornis* under low pH conditions (Putnam et al. 2016) and the freshwater crustacean *Gammarus fossarum* after starvation and thermal stress (Cribiu et al. 2018). It is possible that these higher levels of methylation secure a specific gene expression profile to deal with environmental stress. Across most invertebrates, highly conserved genes or those with housekeeping functions tend to be more heavily methylated (Gavery and Roberts 2014; Keller et al. 2016; Sarda et al. 2012). In contrast, genes that require flexibility and environmental responsiveness tend to have lower methylation levels, allowing for differences in transcriptional regulation (Gavery & Roberts, 2014). However, given that several studies in oysters have found increased methylation associated with higher levels of gene expression, it is possible that increased methylation in stressful conditions allows for higher expression of stress response genes. In this study, a first exposure to hypoxic stress led to reduced variability in methylation among oysters, as shown by tighter clustering of HC and HH replicates on the nMDS. This initial hypoxic stress seems to have had a lasting impact on DNA methylation, influencing the impact of subsequent stressors on DNA methylation patterns, indicative of a carryover effect.

We also observed that methylation responses to hypoxia are not uniform but depend on the timing of stress. Contrary to expectations, oysters that experienced one early exposure to stress (HC) were the most different from all other treatments including oysters that experienced two exposures to hypoxic stress (HH). In fact, both HH and CH oysters only had one significant DMG when compared to oysters that experienced no hypoxic stress (CC), although interestingly, the same gene, MDE1, was differentially methylated in both comparisons. It is possible that our use of whole-body tissue samples might obscure tissue-specific methylation. Alternatively, our conservative thresholds for identifying DMGs may have missed small but critical changes in methylation. Comparing CH and HC oysters resulted in the most DMGs (196) despite both only experiencing a single hypoxic stress exposure demonstrating the importance of timing in shaping the methylome. Oysters that experienced repeated hypoxic stress (HH) had more hypermethylated DMGs (55) compared to oysters where stress was removed (HC; 22 DMGs). Continued stress may necessitate more regulation of specific pathways (resulting in hypermethylation), while a return to normal conditions may allow for greater transcriptional flexibility. Recent exposure to stress in HH and CH conditions appeared to supersede any early life history – we identified no DMGs between HH and CH oysters, indicating that both treatments had similar genomic methylation profiles.

Comparing oysters in a fully factorial design such as in this study adds context to experienced stress and highlights adaptive potentials that may otherwise be missed in other carryover effect studies. For instance, 12 DMGs were identified when comparing CC and HC oysters but it is unclear whether those genes are adaptive. We added context by comparing HH and HC oysters and identified genes (77 DMGs total) that are potentially involved in the adaptive response to hypoxic stress.

### Differentially methylated genes are involved in metabolism, transcription, and stress response

In general, the DMGs we identified were mostly involved in metabolism, transcription, and the stress response, reflecting varied strategies to cope with hypoxic environments. Genes involved in metabolic processes make up the majority (33) of identified DMGs. The duration of hypoxic stress could require different metabolic demands, facilitated by differential methylation of related genes (see section below for connections between methylation and phenotypic change). We see metabolism change in marine invertebrates as a response to stressful environments in order to conserve energy (Hochachka et al. 1996), such as a depression in metabolic rates in Manila clams (*Ruditapes philippinarum*) (Sun et al. 2021) and *C. gigas* exposed to acute hypoxic stress (Haider et al. 2020). Methylation of genes involved in these pathways may be responsible for facilitating changes in energy conservation when oysters experience hypoxic stress. Ten metabolism DMGs were shared in comparisons of CH and HH oysters with HC oysters. Glutamate dehydrogenase (GDH), an enzyme involved in amino acid metabolism, was hypomethylated for HC in both of these pairwise comparisons. This gene is often responsive to environmental changes, such as salinity (Wickes and Morgan II 1976) and pH (Moyes et al. 1985) stress and changes in diet (Li et al. 2025) in bivalves. Methylation of this gene may facilitate differential energy usage to reflect the best strategy in the given environment. Most DMGs were unique to that comparison, suggesting oysters have specific responses within metabolic pathways that differ depending on the timing of hypoxic stress. Some of the identified genes have been previously associated with bivalve stress response. For instance, TRIM71, a gene involved in the protein catabolic process, was hypermethylated in HC oysters when compared to HH oysters. This gene was also highly expressed in *C. gigas* resistant to summer mortality compared to those that are susceptible (Chi et al. 2023), suggesting it may play a role in stress resilience.

We also found many DMGs involved in transcription (10). We noted that transcription-factor E2F3 was hypermethylated in oysters that experienced a single late hypoxic stress (CH) compared to oysters with a single early hypoxic stress exposure (HC). *C. virginica* exposed to air and cold stress downregulated many cell cycle and cell division genes, including E2F3, in a study from (Li et al. 2022). Additionally, Yao et al. (2024) found E2F3 to be a crucial gene responsible for growth regulation in dwarf surf clams (*Mulinia lateralis*). Differential methylation of this gene and others involved in transcription reflect possible fine-tuning of transcriptional control in bivalves.

Thirteen genes related to stress response were differentially methylated in comparisons with HC oysters. Gamma-glutamyl transpeptidase (GGT1) was the one stress related DMG shared in all three pairwise comparisons and was always hypomethylated in HC oysters. This gene codes for an enzyme that synthesizes glutathione, which is a well-known antioxidative agent to combat the production of reactive oxygen species (ROS) in prolonged hypoxic exposure (Hanigan 2014; Margis et al. 2008; Nava et al. 2009). In marine invertebrates, GGT1 is often upregulated in response to environmental stressors, as seen with hypoxic stress in pearl oysters (Luo et al. 2024), salinity stress in Manila clams (Sun et al. 2021), and temperature stress in sea urchins (Liu et al. 2023). Less methylation of GGT1 in HC oysters in all comparisons might enable different hypoxic tolerance capabilities in HC oysters than oysters in the other treatment combinations. Another gene with antioxidative properties, glutathione peroxidase (GPx), was hypermethylated in HC oysters when compared to CH oysters which suggests there are differences in the strategies employed to deal with hypoxic stress, depending on the timing of the stress. A study exposing *C. gigas* to hypoxia found that GPx was upregulated after hypoxic exposure when compared to control oysters (David et al. 2005). More broadly, antioxidant genes seem to play a key role in the hypoxic response in oysters across studies. For example, Sussarellu et al. (2010) showed hypoxia-resistant oysters activated glutathione-S-transferase and other antioxidant enzymes after 20 days of hypoxia. *C. virginica* differentially expressed two alternative splice transcripts of an oxidase gene after oxygenation stress (Liu and Guo 2017). Methylation may influence the inclusion and expression of variable splice forms to plastically respond to environmental changes, like hypoxia.

Only one gene was differentially methylated (hypomethylated for CC) in comparisons between CC and both CH and HH. This gene, MDE1 (methylthioribulose-1-phosphate dehydratase), is involved in the methionine salvage pathway (Albers 2009). Methionine is a precursor metabolite to antioxidative products, like glutathione (Bin et al. 2017). Differential methylation of these antioxidant pathways may alter tolerance to hypoxic stress, depending on the timing of when the oyster experiences the stress. In addition to genes directly related to oxidative stress, heat shock proteins are known to be involved in multiple stress response pathways, allowing cells to react quickly to stressors and often used as a biomarker for stress in bivalves (de Jong et al. 2008; Dimitriadis et al. 2012; Fabbri et al. 2008; Ratkaj et al. 2015; Sørensen et al. 2003). HSP70 was hypermethylated in HC oysters compared to CH oysters. In *C. gigas* (David et al. 2005)*, R. philippinarum* (Nie et al. 2018), and *Mercenaria mercenaria* (Hu et al. 2023), HSP70 was upregulated when exposed to hypoxic stress. Differences in methylation based on prior exposure to hypoxia may be altering expression of heat shock proteins, allowing rapid and adaptive responses to environmental stress.

### Methylation and phenotypic carryover effects

The oysters in the present study are a subset of the oysters used in the experiment from Donelan et al., 2021. In the companion study, we reported that a second exposure to hypoxia, rather than a first exposure, had more of an effect on tissue and shell growth in these oysters. Interestingly, our results show the inverse relationship, where overall differences in DNA methylation are largely driven by an early exposure to hypoxic stress. These different patterns suggest that molecular changes may occur before phenotypic changes like tissue and shell growth are visible, or potentially that the molecular carryover effects observed here are decoupled from the previously documented gross phenotypic carryover effects. Methylation has previously been demonstrated to have the capability to facilitate phenotypic change in bivalves. For example, a study comparing fast and slow growing *C. gigas* lines identified differential methylation of growth-related genes (Tan et al. 2022). Similarly, geoduck clams conditioned to low pH initially experienced changes in their relative shell size and methylome but did not further change upon a second exposure to the same stressor (Putnam et al. 2022). In our study, we do not see strong evidence of DNA methylation acting as the mechanism underlying phenotypic carryover effects. Nonetheless, differences in methylation patterns were based on both the first and second exposure in oysters, indicative of molecular carryover effects. However, at this time, we are limited in understanding how these molecular carryover effects influence phenotypic change in oysters. Future studies will incorporate transcriptomic analysis to further clarify this relationship between DNA methylation and gene expression. Altogether, these findings suggest how an early environmental exposure may encode lasting impacts on the epigenetic landscape of oysters, contributing to our understanding of how bivalves respond to climate change related stressors.

### Limitations

Although we identified DMGs as a result of variations in the timing of hypoxic stress, our results may be confounded by the use of whole-body tissue samples. Many methylation patterns are specific to the tissue type and therefore are potentially obscured in our analyses. Additionally, the use of methylRAD-sequencing in our study resulted in lower coverage than other standard methylation sequencing methods, such as bisulfite sequencing. While this method allowed us to identify differences across treatments, we likely are not capturing the entirety of the methylome. Lastly, the functional role of methylation in most invertebrates, especially oysters, remains unclear. While we observed differences in methylation between treatments, we can only speculate how methylation may influence gene regulation. In the future, studies would benefit from analyzing methylation and gene expression concurrently in tissue-specific contexts to better understand this relationship.

## Data availability statement

Raw sequence data is available at the NCBI Sequence Read Archive under BioProject accession number PRJNA1327452. Associated information for all analyses and supplemental material can be found in the GitHub repository (McDonough 2025; https://github.com/jgmcdonough/CE18_methylRAD_analysis).

## Acknowledgements

We would like to thank Denise Breitburg and Matt Ogburn, Katrina Pagenkopp Lohan, and Greg Ruiz for support and help with the initial study design, Kristina Colacicco and Yamilla Samara Chacon for help performing the experiments, and Groves Dixon and Misha Matz for help troubleshooting the MethylRAD protocol.

